# Drug Washout and Viral Rebound: Modeling HIV Reactivation Under ART Discontinuation

**DOI:** 10.1101/2025.03.22.644757

**Authors:** Mesfin Asfaw Taye

## Abstract

Due to the persistence of latently infected CD4^+^ T cells, achieving a functional cure for HIV-1 remains a significant challenge since the viruses are able to evade immune clearance, which in turn enables post-treatment viral rebound. Because traditional deterministic models assume a constant reactivation rate, they fail to capture the stochastic nature of latency reversal influenced by immune perturbations and ART pharmacokinetics. Thus, in this study, by using a Poisson-driven stochastic framework that incorporates fluctuations in activation rates, we study viral rebound dynamics. Via an exponentially decreasing drug washout model, we accurately quantifies the nonlinear interplay between ART decay and stochastic reactivation, improving the theoretical estimates of post-treatment control. Beyond the introduction of stochasticity, our model establishes a time-dependent viral reactivation framework that integrates periodic and random perturbations in the activation rates. Unlike conventional models that assume uniform (temporally independent reactivation), we show that latency reversal follows structured oscillatory patterns modulated by immune cycles, circadian rhythms, and transient inflammatory episodes. This finding suggests that viral rebound risk is dynamically shaped by immune fluctuations, contrary to the assumption of a constant reactivation probability. We also study the model system by incorporating Gamma-distributed waiting times to account for heterogeneity in reactivation kinetics, which in turn provides a more flexible characterization of reservoir dynamics. We believe that these insights have critical implications for HIV cure strategies. For instance, the shock-and-kill approach relies on latency-reversing agents (LRAs) to reactivate the latent reservoir for immune-mediated clearance. Our findings suggest that periodic immune stimulation could enhance viral clearance, which indicates that synchronizing LRA administration with immune activation cycles may improve therapeutic efficacy. Furthermore, by coupling stochastic reactivation dynamics with ART pharmacokinetics, we identify optimized treatment interruption protocols that potentially delays viral rebound and extending ART-free remission. Moreover, this framework offers a generalizable model for chronic viral infections beyond HIV, including hepatitis B virus (HBV) and cytomegalovirus (CMV) since immune fluctuations and stochastic reactivation play a central role in viral persistence. By expanding theoretical models to incorporate dynamic reactivation rates, immune perturbations, and pharmacokinetic decay, our study refines the predictive modeling of post-treatment control and provides a mathematical foundation for optimizing cure strategies in persistent viral infections. Additionally, we show that the efficacy of latency-reversing interventions, such as the Shock-and-Kill strategy, can be enhanced by synchronizing latency reversal with peak immune activity, improving post-treatment control. Beyond HIV, our framework provides a generalizable model for other persistent viral infections, including hepatitis B virus (HBV) and cytomegalovirus (CMV), offering valuable insights into the interplay between immune dynamics, drug decay, and viral reactivation.

**PACS numbers:** Valid PACS appear here

## I. INTRODUCTION

The advent of antiretroviral therapy (ART) has revolutionized human immunodeficiency virus HIV-1 management since ART is able to successfully suppress viral replication. As a result, ART is able to suppress the propagation of HIV-1 [1–3]. Despite this success, a complete cure of HIV-1 becomes unattainable due to the presence of latently infected CD4_+_ T cells, which act as a long-lived viral reservoir [4–6]. Research shows that these reservoirs are established early during infection. Even in the presence of antiretroviral therapy, the reservoirs continue to flourish as a result of transcriptional silencing and by evading immune clearance [7–9]. When ART is interrupted, reactivation of these latent cells triggers viral rebound [10–12].

In the past, mathematical modeling has served as basic tool to explore the dynamics of highly virulent viruses [14–30] as well as the stochastic nature of HIV latency and reactivation [13, 31, 32]. While early deterministic models assumed uniform reactivation dynamics, recent evidence suggests that latency reversal is a fluctuating and non-uniform process [33–36] due to immune perturbations and environmental factors. This indicates that incorporating a stochastic frameworks that includs reactivation probability distributions, immune fluctuations, and ART kinetics is vital to predict viral rebound timing [37].

Different clinical trials and empirical studies in -human primate (NHPs) show heterogeneity in rebound timing following ART cessation [7, 8]. Because of the interplay between immune activation thresholds, epigenetic regulation, and ART washout kinetics, research shows that early post-ART reactivation events are infrequent compared with later occurrences during ART interruption [9]. This also indicates that viral reactivation is sporadically driven by circadian rhythms, transient inflammation, and immune activation bursts [34, 35].

Because traditional ordinary differential equation (ODE) models fail to capture these complexities, it is vital to consider stochastic differential equation (SDE) models. These models clearly exhibit the effect of fluctuating reactivation rates and immune-mediated viral clearance [12, 38]. Most importantly, frequency-dependent models serve as an important tool because such an approach properly accounts for the variability in latency reversal, immune surveillance, and ART pharmacodynamics [31, 32].

Experimental studies have provided valuable insights into the mechanisms controlling HIV latency. In particular, Jurkat-derived T cell models serves as a tool in exploring chromatin-based transcriptional silencing. On the other hand primary CD4_+_ T cell models offer a more accurate reflection of latency heterogeneity [39, 40]. The effect of latency-reversing agents (LRAs) and immune-based therapies has been studied in non-human primates (NHPs) [41]. Some recent works has also shown that superior efficacy in delaying viral rebound (compared to single-agent strategies) is obtained when a combine treatment using Toll-like receptor (TLR) agonists and broadly neutralizing antibodies (bNAbs) [42].

We want to emphasize that despite significant progress in understanding HIV latency and reactivation, key challenges still persist. Since single-agent latency-reversing agents (LRAs) have modest effect, it is vital to use a combined therapy that couple latency reversal with enhanced immune-mediated clearance [42, 43]. Furthermore, the fluctuating dynamics of viral reactivation also indicates that the timing of therapeutic interventions should to coincide with periods of heightened immune responsiveness [7, 10]. Moreover, due to the the complex interplay between reservoir heterogeneity, stochastic immune activation, and the unpredictable nature of latency reversal, a sustained ART-free remission remains difficult [34, 36, 44]. Addressing these barriers will require innovative strategies that integrate precision medicine approaches with targeted immunological enhancements.

In this paper, we study the effect of time, immune system fluctuations, and ART pharmacokinetics on viral reactivation following ART discontinuation. In this work, we model HIV reactivation as a Poisson-driven stochastic process based on the fact that latency reversal does not occur uniformly but follows oscillatory as well as random patterns influenced by circadian rhythms, transient inflammation, and immune activation bursts. By incorporating an exponentially decreasing drug washout function, we show that the rate of ART decay significantly affects the timing of viral rebound. It is shown that slower drug decay prolongs the activation time, while rapid clearance accelerates the activation time. In order to capture the variability in latency reversal, we consider Gamma-distributed waiting times capture the variability in latency reversal, confirming that early post-ART activation events occur less frequently than later ones. In a recent work, Wu et al. [45] explored the effects of varying activation rates on HIV and SIV reactivation frequency; however, in this work, we present a rigorous mathematical framework to describe the stochastic nature of these activation processes. In this work, we systematically analyze four distinct activation rate scenarios within a time-dependent Poisson process: (i) a constant activation rate, (ii) a sinusoidally varying activation rate, (iii) a randomly fluctuating activation rate, and (iv) an exponentially decreasing activation rate during drug washout. By solving the model system, we explore how cumulative activation functions, expected event counts, and inter-event waiting times behave. This in turn will we provide a comprehensive understanding of how these activation mechanisms shape viral reactivation dynamics.

Furthermore, we explore the effect of sinusoidal and randomly fluctuating activation rates on latency reversal. The immune-driven oscillations play a crucial role in activation timing. This, in turn, leads to clusters of reactivation events during peak immune activity and prolonged latency during the suppression phases. The survival probability as a function of time is examined for different activation models. The results indicate that slow immune-driven fluctuations reduce the reactivation likelihood, whereas strong stochastic perturbations significantly increase rebound risks. Moreover, stochastic reactivation models show that early ART suppression restricts initial viral expansion. When the drug concentration declines, reactivation-driven viral replication follows an exponential trajectory. These findings may indicate that by aligning the timing latency-reversing interventions with immune activation peaks, one can enhance treatment efficacy and improve post-treatment control.

The Shock-and-Kill strategy is particularly affected by the stochastic nature of reactivation and immune variability. We show that effectiveness is significantly improved when the ”shock” phase (where latency-reversing agents (LRAs) induce latent reservoir activation) is synchronized with immune stimulation. This synchronization leads to enhanced viral clearance in the “kill” phase. The ART washout rate is also dictates the efficiency of this approach since slower decay extends the window for immune-mediated clearance while rapid drug clearance increases the risk of uncontrolled viral expansion.

The rest of the paper is structured as follows. In Section II, we discuss the Poisson process of HIV reactivation. In Section III, we consider HIV reactivation with gamma-distributed waiting time. In Section IV, we present mathematical model of viral reactivation. We study the dynamics of virus, latent cells and infected cells. The role of immune response is studied in Section V. In section VI, we present the shock and kill mathematical model. Section VI deals with summery and conclusion.

## II. POISSON PROCESS FOR HIV REACTIVATION WITH DRUG WASHOUT

In the recent work, Wu et al. [45] explored how varying activation rates influence HIV and SIV reactivation. In this work we develop a rigorous model based on a time-dependent Poisson process to analyze four distinct activation rate such as constant, sinusoidal, randomly fluctuating, and exponentially decreasing during drug washout. By deriving exact solutions for the cumulative activation function, we explore how activation variability dictates event clustering, inter-event waiting times and viral dynamics.

To extend this analysis, we introduce a stochastic differential equation (SDE)-based framework that integrates HIV reactivation, immune fluctuations, and antiretroviral therapy (ART) pharmacokinetics. Since our Poisson-driven model incorporates stochastic noise, it helps to understand the probabilistic nature of latency reversal by considering ART washout time. This in turn helps to grasp how drug decay modulates viral resurgence. By incorporating immune perturbations as random fluctuations, we provide a mechanistic understanding of variability in post-treatment control.

Our results reveal that activation rate heterogeneity fundamentally alters viral reactivation timing and this has implications for treatment strategies. In other words, embedding stochasticity into HIV latency models, leads to a refined theoretical framework that informs the optimization of latency-reversing interventions. This approach also provides insights regarding stochastic activation processes in other persistent infections, such as hepatitis B and cytomegalovirus. The approach also has potential applications in immunotherapy and viral eradication strategies.

### A. Stochastic Modeling of Viral Reactivation Dynamics

A fundamental challenge in understanding post-treatment HIV reactivation lies in the stochastic nature of latency reversal and viral rebound following antiretroviral therapy (ART) cessation. In this work work, we address this complexity by developing a Poisson-driven stochastic framework that accounts for immune perturbations, ART pharmacokinetics, and latency-reversing agent (LRA) administration strategies. The probability *P* (*n, t*) of observing *n* reactivated cells at time *t* is given as

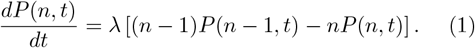

This equation governs transitions between activation states, ensuring that all probability mass remains at *n* = 0 prior to ART discontinuation. At the moment of drug washout (*t* = *t*_*w*_), no reactivation events occurs

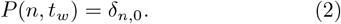

Here *δ*_*n*,0_ denotes the Kronecker delta. After washout, reactivation follows a Poisson process with rate *λ*, yielding a shifted counting process:

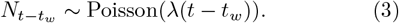

We confirm that the expected number of reactivation events follows:

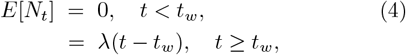

demonstrating a linear increase post-washout.

Our findings also align with Van Dorp et al. [44]. Explicitly solving their stochastic multi-reactivation model, we recover their key result describing viral growth

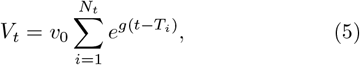

where each reactivated latent cell contributes exponentially to viral replication with rate *g*. The expected viral load post-ART interruption follows:

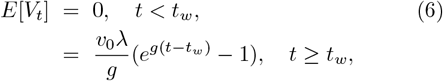

confirming an exponential increase driven by the reactivation rate *λ* and replication dynamics. The variance that exhibits significant stochastic fluctuations

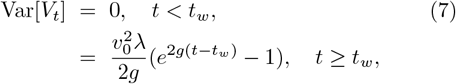

highlights the limitations of deterministic models in capturing latency reversal dynamics. A critical question in treatment interruption studies is the probability of delaying viral rebound. The survival probability *P* (*T > t*) follows an exponential decay:

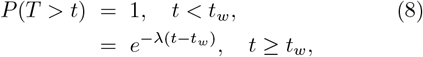

implying that reactivation likelihood increases over time. The waiting time to first activation follows an exponential distribution, with expected time:

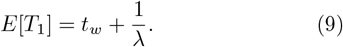

Experimental estimates suggest reactivation rates between 0.17–0.54 per day for HIV and 0.5–2.1 per day for SIV. The expected waiting times rang from 1.9 to 6 days post-washout [7]. More generally, the *k*-th activation event follows a Gamma-distributed waiting time:

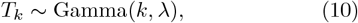

with expected value:

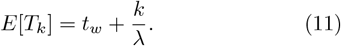

confirming that the expected time to the *k*-th activation increases linearly with the number of activation events. By maintaining reservoir heterogeneity and stochastic fluctuations in latency reversal, this Gamma-distributed waiting-time framework provides a statistical foundation for modeling multiple reactivation events.

Finally, for a constant activation rate *λ*_0_, the survival probability is given as

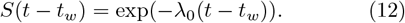

This equation is vital in describing the probability of remaining in latency without reactivation and it also shows the fundamental timescale for viral rebound under continuous stochastic activation. This also helps to gain a basic understanding of HIV latency reversal and treatment interruption strategies by providing a rigorous probabilistic framework for optimizing post-ART control and latency-reversing interventions.

### B. Sinusoidally Modulated Activation Rate

Let us now consider a time-dependent Poisson process with a sinusoidally varying activation rate

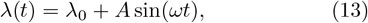

where *λ*_0_, *A* and *ω* denote the baseline activation rate, the amplitude of oscillatory modulation, and denotes the frequency of periodic fluctuations, respectively. Since activation events follow a cyclic pattern, such as circadian rhythms, neuronal firing, and viral reactivation, this formulation is particularly relevant for biological and physical systems.

The cumulative rate function that quantifies the total expected activations up to time *t*, is given by

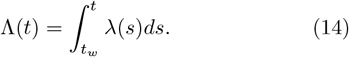

The above equation can be rewriten as

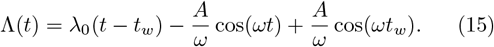

The expected number of activations can be calculated as

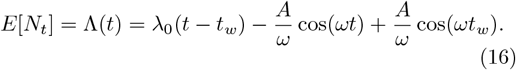

The equation reveals the impact of periodic fluctuations and it shows that activation rates oscillate around their baseline level, contributing to structured clustering of events.

Since inter-event times are inversely proportional to the instantaneous rate, the expected waiting time between consecutive activations is given by:

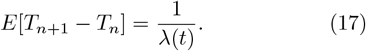

During peak activation phases (sin(*ωt*) ≈ 1), the expected waiting time simplifies to

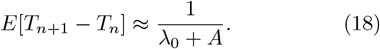

For large oscillatory amplitudes, we get

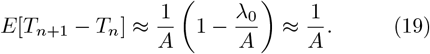

One can clearly see that when *A* is large, activation events cluster during peak phases and this in turn significantly reduces inter-event intervals.

On the other hand, for small oscillations (where *A* ≪ *λ*_0_), one gets

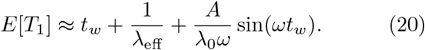

The effective activation rate over an oscillation period is given by

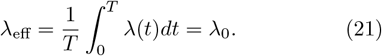

The additional correction term reflects the periodic influences on activation timing and shows deviations from the homogeneous Poisson process.

The expected time for the *k*-th activation can be written as

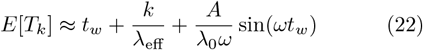

which shows that sinusoidal modulations alter activation event spacing by introducing periodic shifts in inter-event intervals.

To analyze event distributions across an oscillation cycle, let us now compute the cycle-averaged waiting time

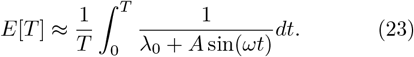

For large oscillations, an asymptotic expansion leads to

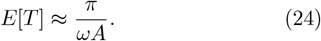

Clearly, the waiting times decrease as *A* increases.

As before, the viral load is given by 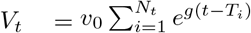. After some algebra, we get the expectation of *V*_*t*_ as 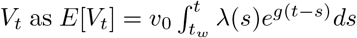. Evaluating this integral, we obtain

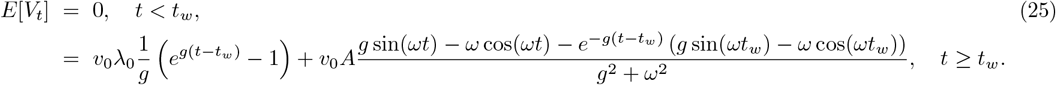

Similarly, the variance of *V*_*t*_ is given by

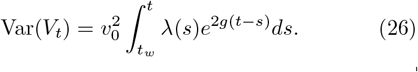

Evaluating this integral, one gets

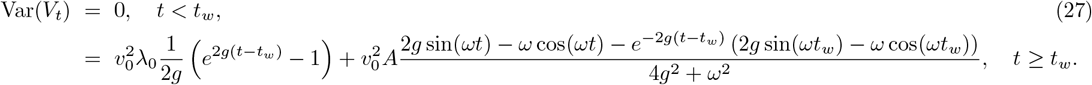

For a periodically varying activation rate, the corresponding survival probability is given by:

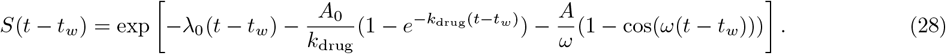

The result obtained in this work also indicates that in the regime of small oscillations (*A* ≪ *λ*_0_), activation events remain relatively uniformly distributed. Inthis case the sinusoidal modulation introduces only minor periodic perturbations. As the result, the effective activation rate remains approximately *λ*_0_ which preserves the statistical properties of a homogeneous Poisson process. In other words, the expected waiting time until activation retains its near-constant behavior, with only slight deviations modulated by the oscillation frequency. In contrast, for large oscillations (*A* ≫ *λ*_0_), the activation rate undergoes substantial fluctuations. This results in a significant clustering of activation events during peak activation phases. The inter-event waiting times become highly non-uniform, with periods of rapid activation alternating with near-suppressed activity. In this regime, the expected inter-event waiting time is well approximated by *E*[*T*] ≈ *π/*(*ωA*). This indicates that activation bursts are primarily dictated by the oscillatory amplitude and frequency. It is vital to note that this transition from near-Poissonian behavior to burst-like activation dynamics has broad implications for biological and physical systems such as neural spike trains, viral reactivation dynamics, and biochemical signaling networks. In neural systems, the clustering of spikes during high-activation phases could influence synaptic plasticity and information encoding. In the field of virology, the intermittent nature of viral reactivation impacts treatment strategies that is aimed at latency reversal. On the other systems such as biochemical networks, oscillatory activation rates may regulate protein synthesis or enzymatic activity. In this case phase-dependent clustering of events could enhance or suppress downstream reactions.

### C. Randomly Fluctuating Activation Rate

In biological systems, activation events are rarely governed by strictly deterministic or periodic mechanisms but are instead modulated by stochastic influences. This is because due to intrinsic noise, cellular variability, and environmental perturbations. Since fluctuations dictate functional outcomes, a randomly fluctuating activation rate provides a crucial framework for capturing biologically relevant activation processes. In biological processes such as neural spike timing, immune cell activation, and viral latency reactivation, activation rates vary dynamically due to molecular noise and external regulatory signals. For neural systems research works indicate that synaptic inputs and membrane potentials fluctuate stochastically which leads to irregular spiking patterns that cannot be fully captured by fixed-rate or purely periodic models. Moreover, since cytokine signaling and immune cell recruitment occurring in bursts driven by infection dynamics and stochastic intercellular interactions, immune responses are highly variable. Latent viruses in HIV or herpes reemerge in a probabilistic manner depending on host immune status, cellular stress, and epigenetic modifications. Gene expression and protein synthesis are also inherently stochastic since transcriptional bursting and translation noise introduce variability in cellular function. Thus incorporating a randomly fluctuating activation rate is vital for a more accurate description of these processes. This also unifies biological activation models by capturing the spectrum from tightly regulated periodic behavior to noise-driven stochasticity.

To account for stochastic variations inherent in biological and physical systems, in this section we consider a Poisson process with a fluctuating activation rate

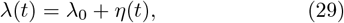

where *λ*_0_ represents the baseline activation rate, and *η*(*t*) is a Gaussian noise process with zero mean and variance:

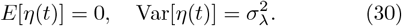

Using the above equation we study the relation of the inherent randomness in activation with neuronal firing, immune responses, and gene expression. For instance, in neuronal networks, synaptic input fluctuations introduce variability in spike timing. The typical baseline firing rates ranges from *λ*_0_ ∼ 5 − 50 Hz. The noise intensity *σ*_*λ*_ can vary depending on synaptic plasticity and external stimuli. In immunology, the activation rate of immune cells responding to an antigen follows stochastic kinetics, with mean activation rates in the range of 0.01 − 1 per hour, modulated by cytokine fluctuations.

Since the expectation of the fluctuating rate is *E*[*λ*(*t*)] = *λ*_0_, we investigate the impact of stochastic fluctuations on the expected waiting time between successive activations. The expected waiting time is given by:

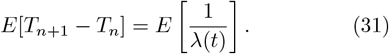

Taylor expanding around *λ*_0_ leads to

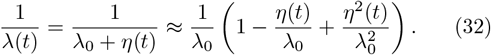

Since *E*[*η*(*t*)] = 0, after some algebra we get

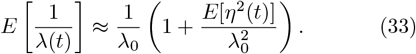

Substituting 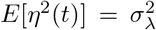, the expected waiting time simplifies to

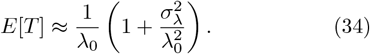

This results indicate that the presence of noise increases the expected waiting time between activation events compared to the deterministic case. The magnitude of this increase depends on the ratio 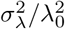 which shows that stronger fluctuations prolong activation intervals.

For large fluctuations, where 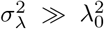, the waiting time is further approximated as:

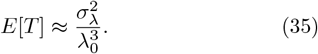

This regime is particularly relevant in biological systems where activation events are dominated by external noise sources, such as gene regulatory networks. In this case the transcription factor binding is probabilistic. Experimental studies suggest that noise-driven gene activation events can occur at rates as low as 10^−4^ per second which shows that significant variability leading to cell-to-cell heterogeneity. The derived expression thus provides a quantitative framework for understanding how fluctuations shape activation timing across diverse stochastic processes.

We further justify this result by using the probability distribution of the cumulative rate function. The probability density function of Λ can be approximated as

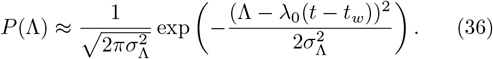

We calculate the firing time as

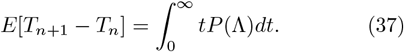

Using integration by parts, one finds

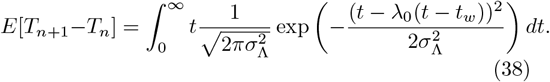

By changing variables and applying Gaussian integral approximations, we derive:

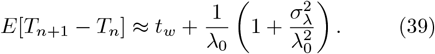

For large fluctuations, this reduces to:

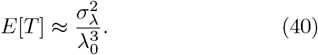

Based on fluctuation regimes, these results can be categorized based on fluctuation regimes. For small fluctuations 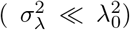, the waiting time remains nearly constant. One can approximate as *E*[*T*] ≈ 1*/λ*_0_. In the case of moderate fluctuations, a quadratic correction emerges. This modifies the waiting time as 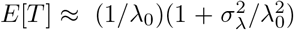. For large fluctuations 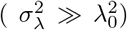, noise effects dominate leading to a waiting time approximation of 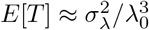.

For exponentially growing viral load 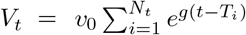, We calculate the expectation of *V*_*t*_ as

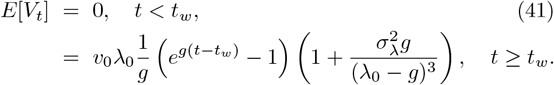

Similarly, the variance of *V*_*t*_ is given by: Var[*V*_*t*_] = 0, *t < t*_*w*_,

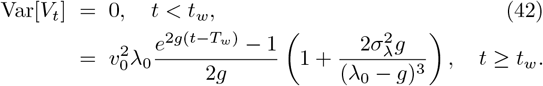

These results highlight the impact of stochastic fluctuations in the activation rate on the expected viral load and its variance. The presence of noise introduces additional corrections, altering the system’s behavior as a function of fluctuation intensity. For small fluctuations where 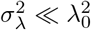, the viral load follows a nearly deterministic trajectory, with deviations governed by the baseline activation rate *λ*_0_. However, for large fluctuations 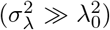, noise dominates, increasing both the expected viral load and its variance. These insights have applications in biological modeling (e.g., gene regulation, neuronal firing), epidemiology (e.g., fluctuating transmission rates in infectious diseases), and stochastic control systems where activation rates exhibit random variability.

The survival probability is given as

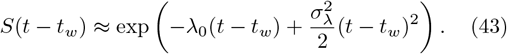

As discussed before, these results have significant implications across various scientific fields. For instance, in neuronal firing cases as well as in gene expression case, the presence of random fluctuations in activation rates can lead to substantial variability in event timing. There fore this fluctuation affects the precision of biological processes that rely on tightly regulated activation sequences. Moreover the time varying activation rates can influence the spread of infectious disease in epidemiological cases. As a result this influence the spread of infectious diseases since noise in transmission rates alters the predicted timing of outbreaks. The insights derived from these equations can be used to optimize stochastic control systems in communication networks. In this case packet transmission times are subject to random delays and understanding the impact of fluctuating activation rates can inform queue management strategies. In a physical system exhibits stochastic resonance—where noise enhances signal detection—can be better characterized through these probabilistic models. These all results indicate that the profound role that noise-induced fluctuations in most dynamic systems. Most importantly, for real biological systems, activation rates are inherently variable. As the rate of fluctuations increases, the system transitions from a nearly deterministic regime to one dominated by stochastic effects.

In Figs. 1 and 2, we plot the probability distribution *P* (Λ) as a function of Λ and *t*, respectively.

**FIG. 1:**
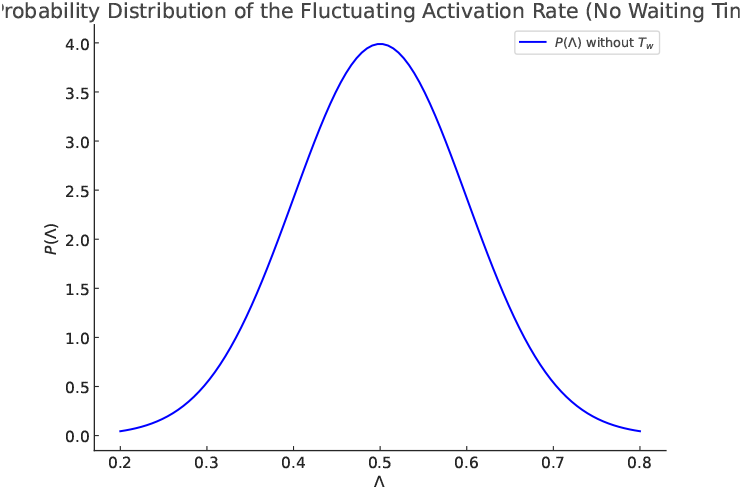
The figure depicts the probability distribution *P* (Λ) of the fluctuating activation rate for a Poisson process with a stochastic rate function *λ*(*t*) = *λ*_0_ + *η*(*t*), where *η*(*t*) is a Gaussian noise process with zero mean and variance 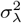. The distribution follows a Gaussian shape centered at the baseline activation rate *λ*_0_ = 0.5 with a standard deviation of *σ*_*λ*_ = 0.1.

**FIG. 2:**
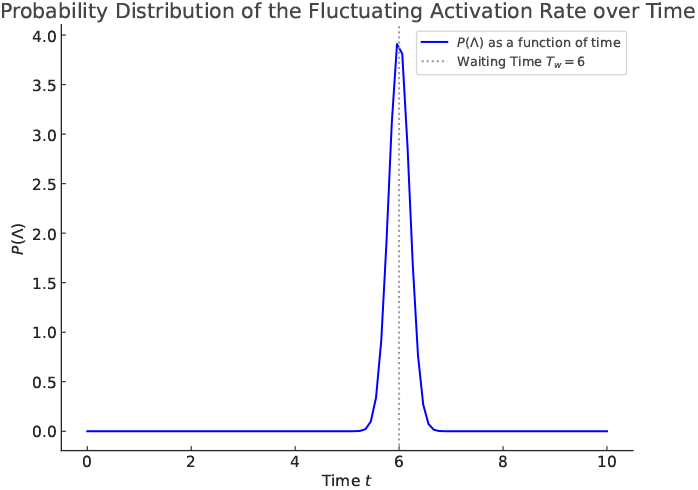
This figure depicts the evolution of the probability distribution *P* (Λ) over time, incorporating a waiting time *T*_*w*_ = 6. The probability density is computed as a function of time, considering the effect of stochastic fluctuations in the activation rate. The vertical dotted line indicates the waiting time *T*_*w*_, before which activation remains suppressed. The parameters used are *λ*_0_ = 0.5 (baseline activation rate) and *σ*_*λ*_ = 0.1 (standard deviation of fluctuations).

### D. Impact of Pharmacokinetic Decay on Latent HIV Activation: A Mathematical Analysis

The activation rate of latent cells following ART interruption is influenced by the pharmacokinetics of drug clearance. To model this effect, we define the time-dependent activation rate as

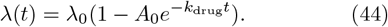

Here *λ*_0_, *A*_0_ and *k*_drug_ denote the baseline activation rate in the absence of ART, the initial suppression effect exerted by the drug, and the drug elimination rate, respectively. At *t* = 0, the activation rate is fully suppressed (*λ*(0) = 0) when *A*_0_ = 1. As the drug concentration declines exponentially over time, the activation rate gradually increases, As time progresses, it approaches its maximum value *λ*_0_ as *t* → ∞.

The rate at which reactivation occurs depends critically on the parameter *k*_drug_. When *k*_drug_ is small, drug clearance becomes slow. This leads to a prolonged suppression of viral reactivation. On the other hand for large *k*_drug_, drug elimination becomes fast. This in turn accelerates the resurgence of viral replication. This findings also reflect that both *λ*_0_ and *k*_drug_ plays a crucial role in determining the timing and extent of viral resurgence by providing a quantitative framework for studying post-treatment control and optimizing therapeutic strategies. Next to model the decay of antiretroviral therapy (ART) concentration over time, let us assume an exponentially decreasing function

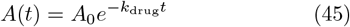

where *A*_0_ and *k*_drug_ denote the initial drug concentration, and the drug elimination rate, respectively. We show that as ART concentration declines, viral replication and latent cell activation gradually resume.

To analyze the impact of pharmacokinetic decay on HIV latent cell activation, we model the cumulative hazard function as 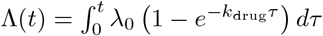. After some algebra, we get 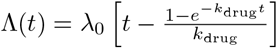.

The expected activation time is given by 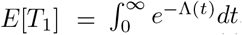. For **small** *k*_drug_, expanding the exponential term in Λ(*t*) and solving perturbatively,

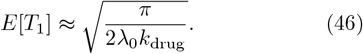

This clearly depicts that delayed activation due to prolonged drug presence. For **large** *k*_drug_, rapid drug clearance leads to

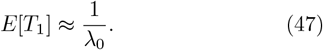

This also indicates that sufficiently fast drug decay, activation time converges to its Poissonian limit.

For moderate *k*_drug_, retaining dominant correction terms, one gets

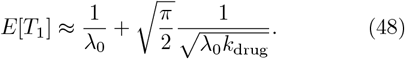

This result highlights the diminishing influence of drug clearance rate beyond a certain threshold. This analysis shows that while drug presence initially delays activation, its long-term impact diminishes as clearance accelerates. The fundamental activation timescale remains governed by *λ*_0_. This also implies that pharmacokinetic decay must be considered in therapeutic strategies but does not fundamentally alter the stochastic nature of latency reversal.

After some algebra, the expectation of *V*_*t*_ is given by

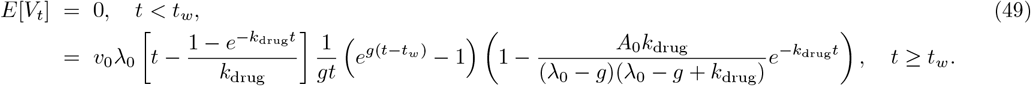

The variance of *V*_*t*_ is caculated as

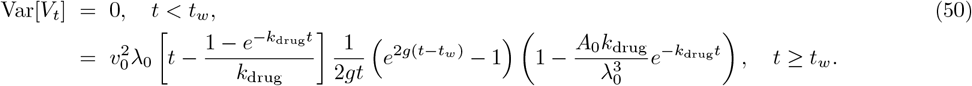

All of these results highlight the role of drug elimination dynamics in shaping viral reactivation timing. Slow clearance extends latency. On the other hand rapid decay restores activation to its baseline level. This analysis provides a quantitative framework for optimizing post-treatment control strategies and understanding latent cell reactivation dynamics in HIV treatment protocols.

The survival probability after some algebra is given by

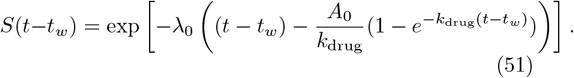

In Fig. 3, the survival probability *S*(*t*) is plotted as a function of time *t* for different activation rate models with a waiting time *T*_*w*_ = 2. Before the waiting time *T*_*w*_, *S*(*t*) = 1, since activation has not started. After the activation time *T*_*w*_, *S*(*t*) decreases based on the activation model: constant rate, drug washout, sinusoidal activation, and stochastic fluctuations. The stochastic model introduces small perturbations around a baseline rate. The vertical dotted line marks *T*_*w*_. Parameters: 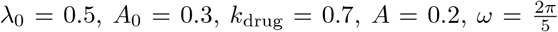, and *η*(*t*) ∼ 𝒩 (0, 0.1). Fig. 4 we plot the survival probability *S*(*t*) is plotted as a function of time *t* for sinusoidal activation with varying amplitude *A*. Larger *A* increases oscillations, affecting survival probability. In the figure other parameters are fixed as 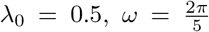, and *A* ∈ {0.1, 0.3, 0.5}.

**FIG. 3:**
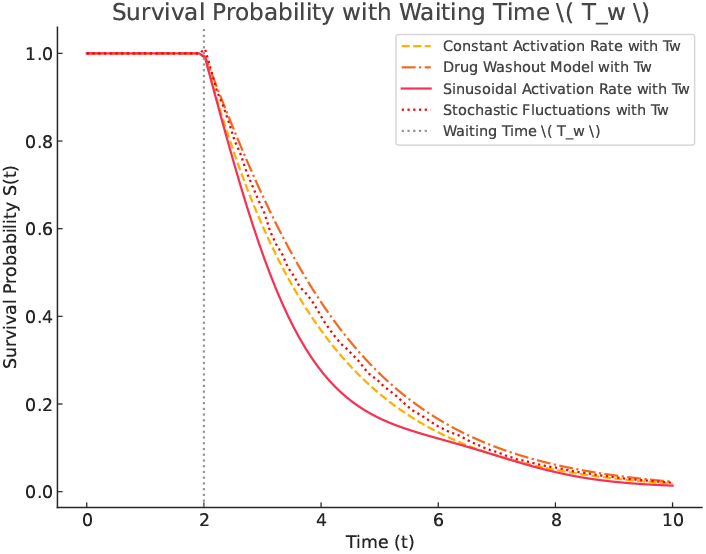
The survival probability *S*(*t*) as a function of time under different activation rate models, incorporating a waiting time *T*_*w*_ = 2. Before *T*_*w*_, the survival probability remains unity, as activation has not commenced. After *T*_*w*_, the probability decreases according to different models: a constant activation rate, a drug washout model, a sinusoidal activation model, and a stochastic fluctuation model. The stochastic model includes small perturbations around a baseline activation rate. The vertical dotted line indicates the waiting time *T*_*w*_. The parameters used are *λ*_0_ = 0.5 (baseline activation rate), *A*_0_ = 0.3 (drug effect strength), *k*_drug_ = 0.7 (drug decay rate), *A* = 0.2 (sinusoidal amplitude), 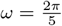 (sinusoidal frequency, corresponding to a period of 5), and stochastic fluctuations modeled as *η*(*t*) ∼ 𝒩 (0, 0.1) with mean zero and standard deviation of 0.1.

**FIG. 4:**
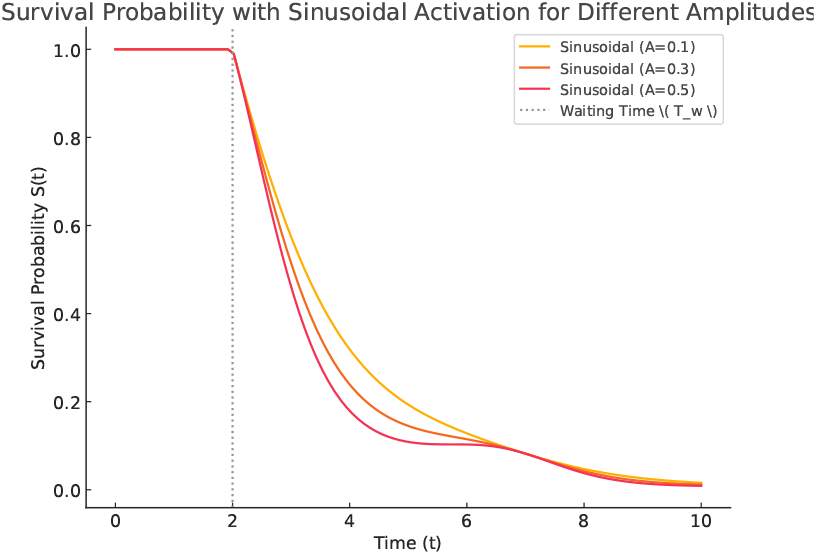
The survival probability *S*(*t*) as a function of time *t* for a sinusoidal activation rate model for different *A*. Here we fix *T*_*w*_ = 2. The sinusoidal activation rate introduces periodic fluctuations in activation probability which affects survival dynamics. As shown in the figure, larger amplitudes lead to stronger oscillations which may result in more pronounced variations in survival probability over time. The vertical dotted line represents the waiting time *T*_*w*_. We fixed other parameters as *λ*_0_ = 0.5 (baseline activation rate),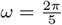 (sinu-soidal frequency, corresponding to a period of 5), and three different amplitudes *A* = [0.1, 0.3, 0.5].

## III. MODELING HIV REACTIVATION WITH A GAMMA-DISTRIBUTED WAITING TIME

Mathematical models of HIV latency traditionally assume that viral reactivation follows an exponentially distributed waiting time, leading to a Poisson-like process. However, biological evidence suggests significant heterogeneity in reactivation times across infected cells. This also implies that indicating that the Poissonian assumption may not fully capture the underlying dynamics.In this study, we derive the analytical expressions for both the expected activation time and the expected number of activations under the assumption that the HIV reactivation process follows a Gamma-distributed waiting time. By using such approach we extend the classical Poisson model to a more flexible and biologically realistic framework than can describe viral latency reversal. Standard HIV reactivation models assume exponential waiting times with a constant hazard rate for viral activation. However, empirical studies indicate substantial variation in the duration of HIV latency across individuals and within cell populations. This heterogeneity suggests that a Gamma-distributed waiting time may provide a more accurate representation of the stochastic process that governs viral reactivation. In this case we derive key mathematical properties of HIV latency reversal are derived by assuming that latent cell activation times follow a Gamma distribution. Specifically, the expected activation time, denoted *E*[*T*], quantifies the mean duration a latent cell remains dormant before reactivation. Furthermore, the expected number of activations, denoted *E*[*N*_*t*_], characterizes the cumulative number of reactivations up to a given time *t*.By developing these properties analytically, the classical Poisson-based models are extended to a more rigorous framework that accounts for heterogeneous activation times.

Let now assume that the waiting time until HIV reactivation follows a Gamma distribution with shape parameter *k* and rate parameter *θ*. The probability density function can be written as

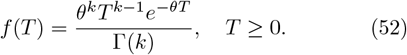

The cumulative distribution function is give by

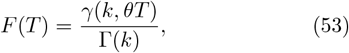

where *γ*(*k, θT*) denotes the lower incomplete Gamma function. The expected activation time has a form

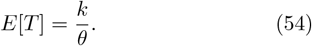

For *k* = 1, the Gamma distribution reduces to the exponential distribution which recovers the Poisson process,

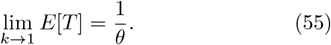

In this section, we consider the previous model system where the drug concentration exponentially decays. The expected activation time *E*[*T*_1_] depends on the pharmacokinetic decay rate *k*_drug_. For small *k*_drug_, one gets

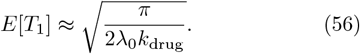

For moderate *k*_drug_, retaining dominant correction terms,

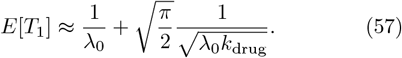

For large *k*_drug_ (where drug clearance is rapid), we get

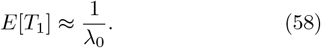

For *k >* 1, activation times are more spread out and this increases the variability in latent cell reactivation. The expected number of reactivations up to time *t* is given as

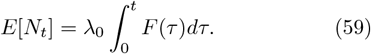

After some algebra, one gets

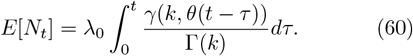

For large *t*, the lower incomplete Gamma function satisfies the asymptotic relation

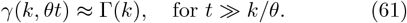

Approximating for large *t*,

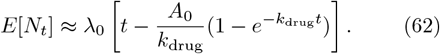

As *k* → 1, the Poisson model is recovered, confirming that the Gamma process generalizes the Poisson case.

In the next section, we consider a deterministic model and explore the interaction of latent cells, free virus, and infected cells. We introduce a pharmacokinetic model to study the dynamics of viral reactivation. The also study the impact of immune response that plays a critical role in controlling viral reactivation. Furthermore, we examine the “Shock-and-Kill” strategy, a promising approach for eradicating latent HIV reservoirs by strategically coordinating latency reversal agents (LRAs) with antiretroviral therapy (ART).

## IV. DETERMINISTIC MATHEMATICAL MODELING OF VIRAL REACTIVATION: INVESTIGATING THE DYNAMICS OF LATENT CELLS, FREE VIRUS, AND INFECTED CELLS WITH AND WITHOUT HOST INTERACTION

### A. The role of latent cells, infected cells, and free virus

Let us now explore the dynamics of latent cells (in the absence of host cells), infected cells, and free virus. In order to investigate the model system, we write a mathematical equation that govern the dynamics of latent cell activation, infection dynamics, and viral replication. The dynamics of the latent cell population is governed by

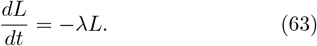

After some algebra, we get

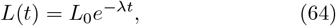

where *L*_0_ represents the initial latent reservoir size. This indicates that in the absence of reactivation or external stimuli, the latent cell population declines over time due to intrinsic cell death or removal mechanisms. The transition from latent to actively infected cells occurs at rate *λL*, while infected cells are removed at rate *δI*. The infected cells evolve as

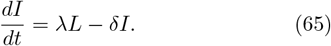

After some algebraic manipulation, one gets

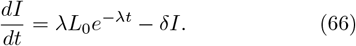

The infected cells depend on time as

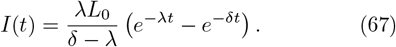

Exploiting the above equation one can see that the infected population initially increases as latent cells transition into actively infected cells. As time progresses, the infected cells either die or are cleared. *I*(*t*) reaches a peak before gradually declining. Moreover, the virus is produced at rate *pI* from infected cells and is cleared at rate *cV*

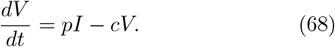

after substituting *I*(*t*), the above equation is simplfied as

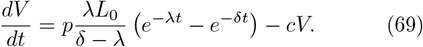

Solving the above equation, we get

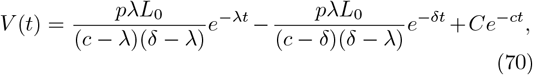

where *C* is determined by initial conditions. From the above equation, one can see that the viral load initially rises due to the production of virions by infected cells. As time progresses, viral load begin to decline. The longterm behavior of the infection can be affected by the competition between viral replication and immune clearance. If clearance rate is sufficiently strong, the virus may be eliminated; otherwise, it may persist at low levels.

The temporal evolution of infection is influenced by physiological variations in key parameters governing latent cell activation, viral production, and clearance rates. In Fig. 5, we show the time evolution of infected cells, *I*(*t*), for two distinct parameter sets. For first set, we use fixed parameters *λ* = 0.2, *δ* = 0.6, *L*_0_ = 5 × 10^4^, *p* = 120, *c* = 2.8, and *C* = 800. for the second set, we fix *λ* = 0.05, *δ* = 0.4, *L*_0_ = 2 × 10^5^, *p* = 80, *c* = 3.5, and *C* = 1500. The figure also depicts that the infected cells increase and peak at a certain time before declining due to clearance mechanisms. In Fig. 6, we plot the viral load *V* (*t*) as a function of time *t*. Depending on viral production and clearance rates, the viral trajectories varies and this may affect infection persistence and decline. As time progresses, viral levels increase and peaks before decreasing. The first parameter set leads to a faster clearance of infected cells and lower viral persistence, whereas the second set exhibits prolonged viral presence, emphasizing the role of physiological variability in infection outcomes.

**FIG. 5:**
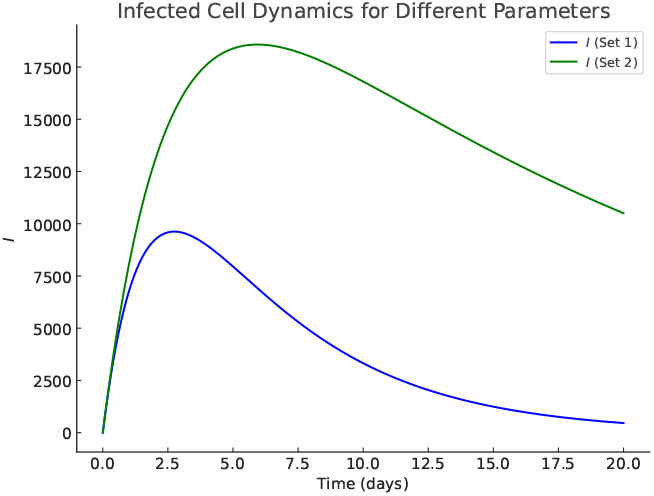
Time evolution of infected cells, *I*(*t*), for two different parameter sets representing physiological variations. The first set uses *λ* = 0.2, *δ* = 0.6, *L*_0_ = 5 × 10^4^, *p* = 120, *c* = 2.8, and *C* = 800. The second set uses *λ* = 0.05, *δ* = 0.4, *L*_0_ = 2 × 10^5^, *p* = 80, *c* = 3.5, and *C* = 1500. These parameters influence the activation and clearance rates, leading to distinct infection dynamics over time.

**FIG. 6:**
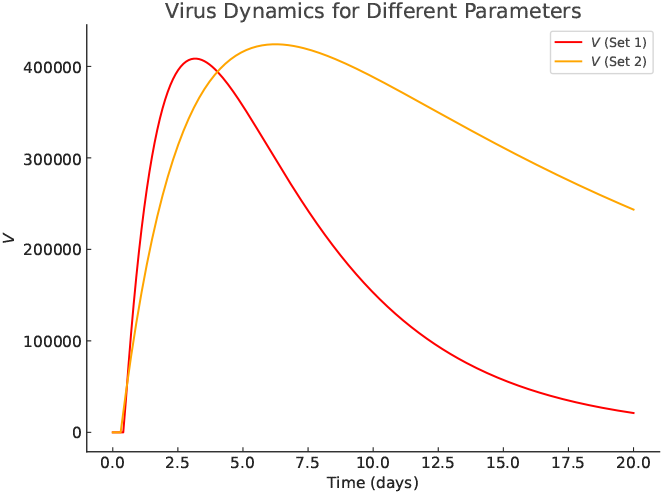
Time evolution of the viral load, *V* (*t*), under two different physiological parameter sets. The first set uses *λ* = 0.2, *δ* = 0.6, *L*_0_ = 5 × 10^4^, *p* = 120, *c* = 2.8, and *C* = 800. The second set uses *λ* = 0.05, *δ* = 0.4, *L*_0_ = 2 × 10^5^, *p* = 80, *c* = 3.5, and *C* = 1500. The variation in production and clearance rates results in distinct viral dynamics, influencing infection persistence and decline.

### B. Viral Reactivation in the Presence of host Cells

Let us not consider the dynamics of the system in presence of host cells. Prior to drug washout, antiretroviral therapy (ART) effectively suppresses viral replication. This prevents new infections and latent cell activation. The latent cell population remains constant since ART prevents activation

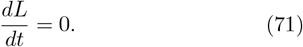

Similarly, the population of infected cells is stabilized as ART inhibits viral replication and the formation of new infections

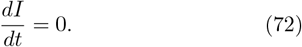

The target cell population follows natural regeneration and decay in the absence of significant viral interactions

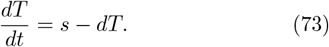

The viral load remains at a suppressed level since ART prevents active replication:

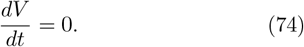

Let us now consider the dynamics after drug washout (*t* ≥ *t*_*w*_) (following drug washout). The suppression of viral replication leads to viral rebound. As a result, the latent cell become activated and more host cells become infected. To account for ART decay over time, we introduce a pharmacokinetic model for drug clearance as

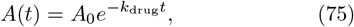

where *A*_0_, *k*_drug_ and *λ* denote the initial drug concentration, the drug elimination rate and the activation rate of latent cells, respectively. The activation rate depends on ART as

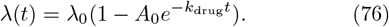

Thus, the latent cell population decreases due to activation but is replenished by new latently infected cells formed through viral interactions

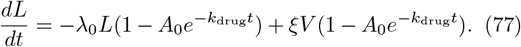

The infected cell population increases as latent cells reactivate and new target cells become infected. As a result

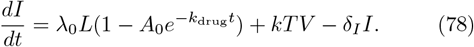

Target cells continue to regenerate naturally but are depleted due to infection by the virus. Hence

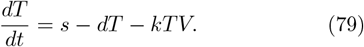

The viral load increases as infected cells produce new virions, while clearance occurs at a natural rate. Therefore, the dynamics of the virus is goverend by

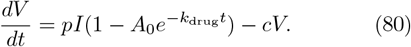

Before drug washout, ART maintains viral suppression by preventing latent cell activation and new infections. As a result the system become stable. However, after ART cessation, the drug concentration decreases over time and this progressively allows viral replication to resume. This leads to latent cell activation, increased infection rates, and viral rebound. The presence of ART decay via 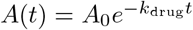 leads to a more accurate representation of post-treatment dynamics. Since the decay highlights the gradual transition from viral suppression to active infection. The presence of time-dependent *λ*(*t*) accounts for varying activation rates of latent cells based on ART availability, improving the model’s biological relevance.

Before drug washout, antiretroviral therapy (ART) effectively suppresses viral replication since it prevents new infections as well as latent cell activation. At equilibrium, we get

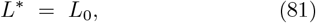

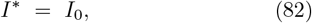

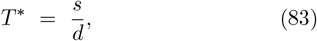

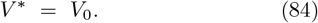

From these equations, one can see that the stabilizing effect of ART. The latent cells remains without activation. The infected cells and viral load remain at negligible levels. The stability of this state underscores the necessity of continuous ART adherence any drug interruption may disrupt this balance which leads to viral rebound.

After ART is stopped, we write relation at equilibrium as

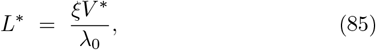

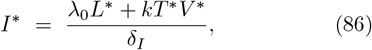

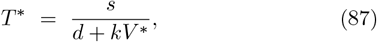

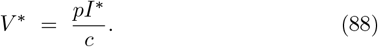

These equilibrium conditions provide key insights into the post-treatment dynamics of HIV. The equations depict that the viral load increases following ART cessation, This also leads to a chain reaction of latent cell activation and immune system destabilization.

The speed of viral rebound depends on the rate at which 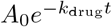 declines showing the importance of understanding ART pharmacokinetics when designing treatment strategies. These results suggest that abrupt ART discontinuation may accelerate disease progression, whereas gradual withdrawal or intermittent ART cycling could mitigate the severity of reactivation by modulating the availability of susceptible target cells.. The above equtions also suggest that the effectiveness of latency-reversing agents (LRAs) depending on the timing of their administration relative to viral load fluctuations. Specifically, LRA interventions may achieve greater reductions in the latent reservoir when they are administered at points where *V* ^∗^ remains low so that the risk of reseeding new latent cells become minimized. Moreover, the dependence of viral and infected cell equilibrium states on the ART decay function, indicates that structured ART interruptions may be optimized to delay rebound while minimizing cumulative drug exposure.

Overall, the equilibrium analysis of the system before and after drug washout is important in order to understand the fundamental mechanisms driving HIV persistence and reactivation. Prior to ART cessation, the system remains in a stable, suppressed state, ensuring viral control. However, following drug elimination, the viral reservoir reactivates. this lead to an active infection state driven by viral replication, target cell depletion, and latency reversal. The incorporation of a time-dependent ART decay model provides a more biologically relevant representation of treatment dynamics, revealing key insights into therapeutic interventions aimed at minimizing rebound and enhancing viral suppression strategies.

Let us now explore the model system. In Fig. 7, we plot the time evolution of latent cells, *L*(*t*) as function of time for different drug decay rates, *k*_*d*_ = 0.001, 0.01, 0.1, 1.0. The results illustrate that as the drug washes out more slowly, the relaxation time for latent cells extends by delaying activation. Fig. 8 depicts *V* (*t*) as a function of time *t* for the same drug decay rates as Fig.7. The rate at which the drug decays significantly impacts viral clearance and persistence. A prolonged drug washout results in a substantial decrease in viral load, whereas faster decay leads to more rapid viral resurgence. The parameters used in both simulations are *s* = 100000, *λ* = 1 × 10^−6^, *d* = 0.1, *h* = 0.1, *e* = 1 × 10^−10^, *d*_1_ = 1.0, *p*_1_ = 10^5^, *c*_1_ = 29, *A*_0_ = 0.3, and *T*_0_ = 10^5^.

**FIG. 7:**
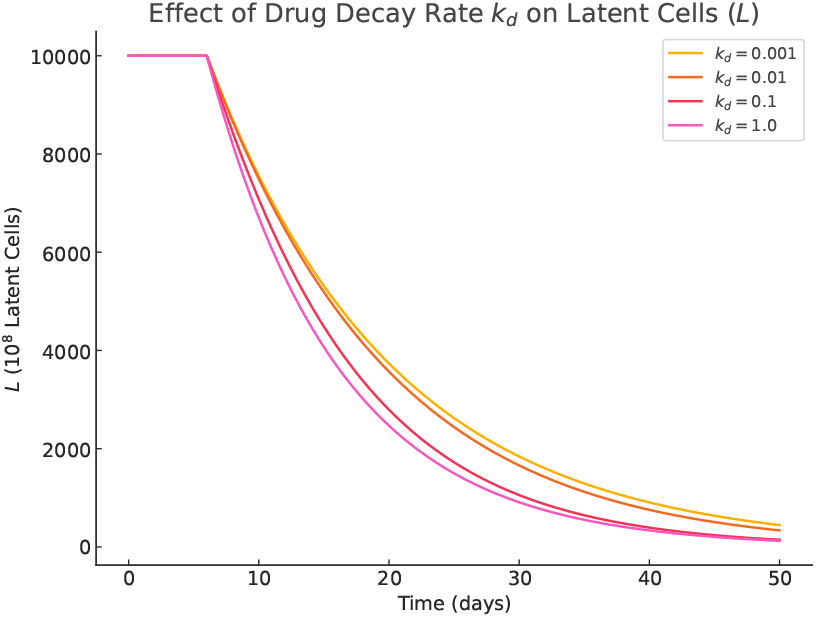
Time evolution of latent cells, *L*(*t*), for different drug decay rates, *k*_*d*_ = 0.001, 0.01, 0.1, 1.0. The latent cell dynamics are influenced by the rate of drug decay, affecting the activation of the latent reservoir. The parameters used in the model are: *s* = 100000, *λ* = 1 × 10^−6^, *d* = 0.1, *h* = 0.1, *e* = 1 × 10^−10^, *d*_1_ = 1.0, *p*_1_ = 10^5^, *c*_1_ = 29, *A*_0_ = 0.3, and *T*_0_ = 10^5^.

**FIG. 8:**
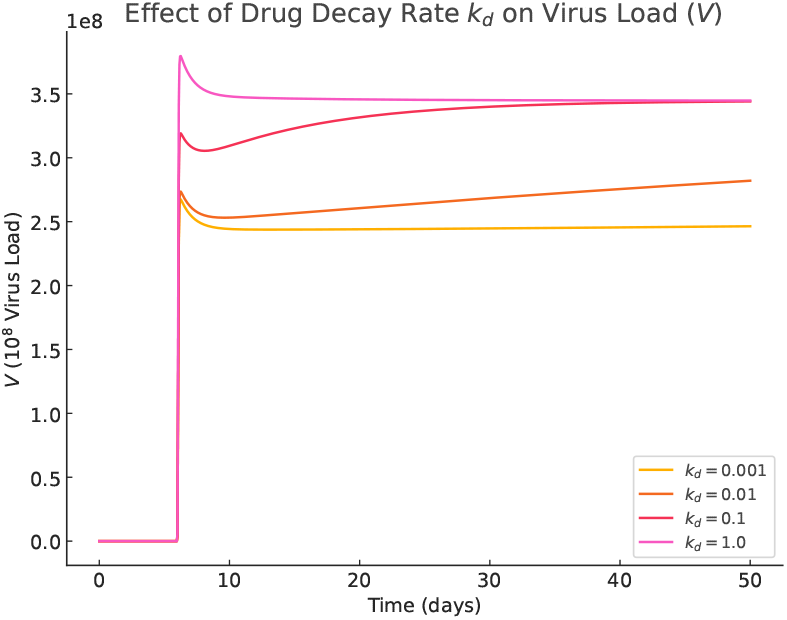
Time evolution of the virus load, *V* (*t*), for different drug decay rates, *k*_*d*_ = 0.001, 0.01, 0.1, 1.0. The viral load is significantly affected by the rate at which the drug decays, leading to different viral clearance or persistence dynamics. The parameters used in the model are: *s* = 100000, *λ* = 1 × 10^−6^, *d* = 0.1, *h* = 0.1, *e* = 1 × 10^−10^, *d*_1_ = 1.0, *p*_1_ = 10^5^, *c*_1_ = 29, *A*_0_ = 0.3, and *T*_0_ = 10^5^.

To ensure biological realism, the model employs physiologically relevant parameter values derived from experimental and clinical studies. The following table provides a comprehensive overview of the key parameters This parameter table reflects established physiological ranges while incorporating variability to account for patient heterogeneity. T

**TABLE I:** Physiological Parameters Used in the Model.

| Parameter | Symbol | Value Range |
| --- | --- | --- |
| Latent cell reactivation rate | $\lambda$ | 0.01 – 0.1 per day |
| Latent cell formation rate | $\xi$ | Variable |
| Latent cell decay rate | $\delta_L$ | $10^{-4} - 10^{-2}$ per day |
| Infected cell death rate | $\delta_I$ | 0.5 – 1.0 per day |
| Target cell regeneration rate | $s$ | $10^3 - 10^5$ cells/μL/day |
| Target cell decay rate | $d$ | $10^{-2} - 10^{-1}$ per day |
| Viral infection rate | $k$ | $10^{-8} - 10^{-6}$ mL/virion/day |
| Virus production rate | $p$ | $10^2 - 10^4$ virions/cell/day |
| Virus clearance rate | $c$ | 10 – 30 per day |
| Immune stimulation rate | $\alpha$ | $10^{-2} - 10^{-1}$ per day |
| Effector cell decay rate | $\mu$ | $10^{-2} - 10^{-1}$ per day |

In Fig. 9, we present *L*(*t*) as a function of time *t* for different initial latent cell counts, *L*_0_ = 1 × 10^3^, 1 × 10^4^, 1 × 10^5^, 1 × 10^6^. As depicted in the figure, the time evolution of latent cells is influenced by the initial size of the latent reservoir. The results indicate that larger initial latent reservoirs lead to prolonged decay times since large latent cells reservoir require more time to clear. Fig. 10 shows the time evolution of infected cells, *I*(*t*) for the same initial latent cell counts as Fig. 9. The dynamics of infected cells depend on both the initial latent reservoir and the activation rate, this also determines the transition of latent cells into the actively infected state. Higher initial latent cell concentrations result in increased infected cell populations by sustaining a higher viral load over time. Fig. 11 depicts the plot *V* (*t*) as a function of *t* for the same initial latent cell counts as Fig. 9. The figure shows that the viral load is strongly dictated by the initial latent reservoir size. Larger *L*_0_ values results in greater viral replication and extended persistence. The parameters used in all simulations are *s* = 1 × 10^5^, *λ* = 1 × 10^−6^, *d* = 0.1, *h* = 0.1, *e* = 1 × 10^−10^, *d*_1_ = 1.0, *p*_1_ = 10^5^, *c*_1_ = 29, *A*_0_ = 0.3, *k*_*d*_ = 0.1, and *T*_0_ = 10^5^.

**FIG. 9:**
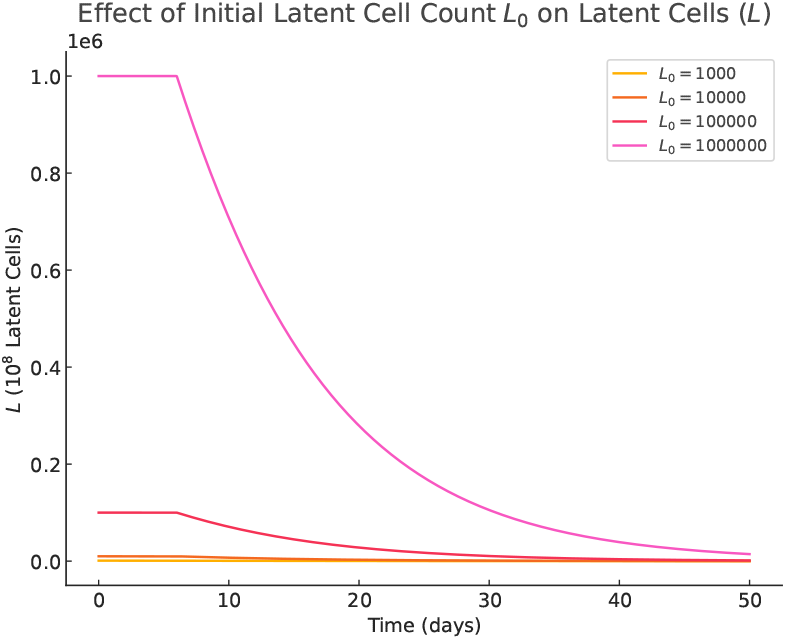
Time evolution of latent cells, *L*(*t*), for different initial latent cell counts, *L*_0_ = 1 × 10^3^, 1 × 10^4^, 1 × 10^5^, 1 × 10^6^. The latent cell dynamics are influenced by the initial size of the latent reservoir. The parameters used in the model are: *s* = 100000, *λ* = 1 × 10^−6^, *d* = 0.1, *h* = 0.1, *e* = 1 × 10^−10^, *d*_1_ = 1.0, *p*_1_ = 10^5^, *c*_1_ = 29, *A*_0_ = 0.3, *k*_*d*_ = 0.1, and *T*_0_ = 10^5^.

**FIG. 10:**
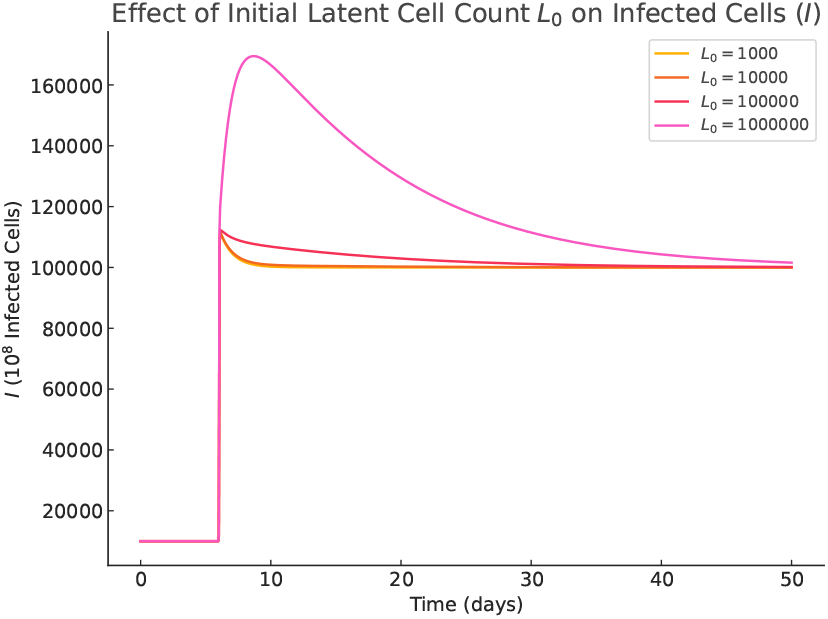
Time evolution of infected cells, *I*(*t*), for different initial latent cell counts, *L*_0_ = 1 × 10^3^, 1 × 10^4^, 1 × 10^5^, 1 × 10^6^. The dynamics of infected cells are impacted by the initial latent cell pool, as well as the rate of activation and infection. The parameters used in the model are: *s* = 100000, *λ* = 1 × 10^−6^, *d* = 0.1, *h* = 0.1, *e* = 1 × 10^−10^, *d*_1_ = 1.0, *p*_1_ = 10^5^, *c*_1_ = 29, *A*_0_ = 0.3, *k*_*d*_ = 0.1, and *T*_0_ = 10^5^.

**FIG. 11:**
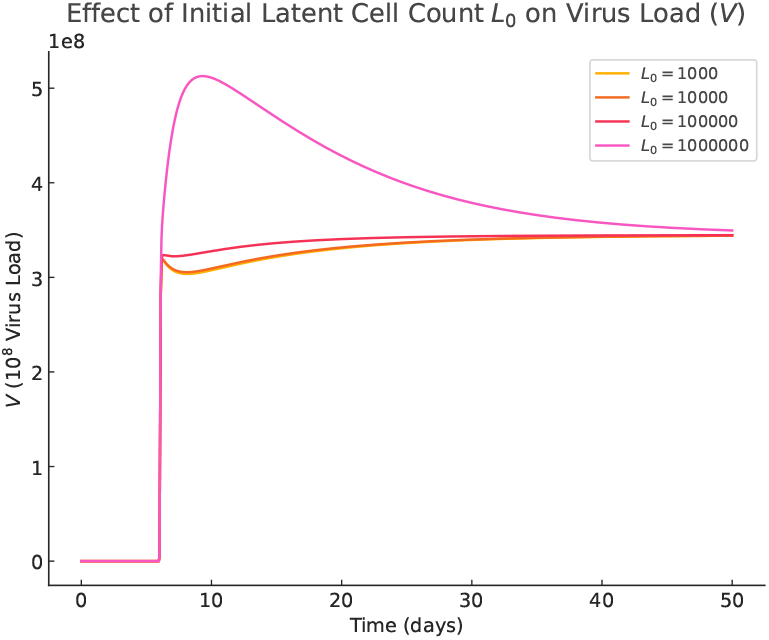
Time evolution of the virus load, *V* (*t*), for different initial latent cell counts, *L*_0_ = 1 × 10^3^, 1 × 10^4^, 1 × 10^5^, 1 × 10^6^. The virus load is influenced by the size of the initial latent reservoir, with higher initial *L*_0_ values resulting in larger viral replication and persistence. The parameters used in the model are: *s* = 1 × 10^5^, *λ* = 1 × 10^−6^, *d* = 0.1, *h* = 0.1, *e* = 1 × 10^−10^, *d*_1_ = 1.0, *p*_1_ = 10^5^, *c*_1_ = 29, *A*_0_ = 0.3, *k*_*d*_ = 0.1, and *T*_0_ = 10^5^.

This mathematical model helps us understand how HIV, remain dormant and reactivate over time. Infection with the virus such as HIV are challenging to eliminate because some infected cells stay hidden in the body and occasionally become active again. As a result antiretroviral therapy (ART) alone cannot completely cure the infection. ART may suppress the viral load but it does not remove these hidden reservoirs. Our model also shows how viruses can evade the immune system by allowing them to persist for long periods despite the body’s defenses. From a treatment perspective, studying the balance between dormant and active virus states can help improve therapies. By understanding when and why reactivation happens, researchers can develop better strategies to reduce hidden infections and keep the virus under control.

## V. IMMUNE RESPONSE AND EFFECTOR CELL DYNAMICS

Let us now consider the effect of the immune response. The viral reactivation is controlled by the immune response by removing infected cells as well as suppressing viral load. Effector cells (*E*) are introduced into the model to account for immune-mediated clearance, governed by

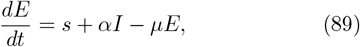

where *s* represents baseline immune activation, *αI* captures immune stimulation by infected cells, and *µE* describes natural effector cell decay.

The effector and infected are related via

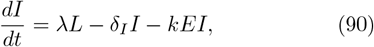

where *kEI* represents immune-mediated clearance. A strong immune response (*E* ≫ 0) accelerates infected cell removal, while a weak response allows viral persistence. Latent cells (*L*) serve as a long-term viral reservoir, with dynamics given by

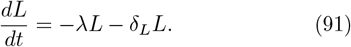

Netx, we investigate the early time approximation of drug washout. Initially, when immune activation is weak (*E* ≈ 0), the term *kEI* is negligible, reducing the infected cell equation to

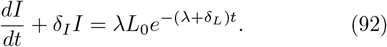

After some algebra one gets

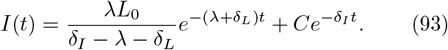

As drug washout progresses, *k*_*d*_ decreases, leading to a longer decay time for *I*(*t*). This aligns with experimental findings where slower drug decay prolongs viral suppression.

The viral load follows

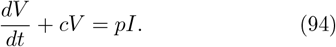

Applying the integrating factor and substituting *I*(*t*), we obtain

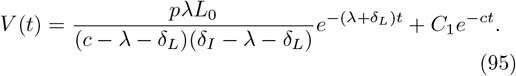

This result indicates that prolonged drug efficacy (*k*_*d*_ ≪ 1) results in a slower decay of viral load.

Setting all derivatives to zero, the system’s equilibrium is determined by

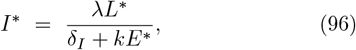

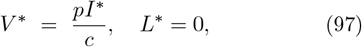

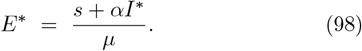

As it can be seen from the above equation, ART suppresses viral replication and latent reservoirs become stabilize. However, if ART is cycled as ON and OFF, latent cells deplete as periodic viral bursts drive immune clearance. This aligns with the “shock and kill” strategy, where latency-reversing agents aim to exhaust the reservoir through controlled immune activation. Note that time-dependent solutions of *I*(*t*), *V* (*t*), and *E*(*t*) require numerical methods such as Runge-Kutta integration.

## VI. OPTIMIZED SHOCK-AND-KILL STRATEGY WITH ART CYCLING AND LATENCY REVERSAL AGENTS

The ”Shock-and-Kill” strategy is one of the most important strategy that helps to eliminate latent HIV by combining latency reversal agents (LRAs) with antiretroviral therapy (ART). LRAs activate dormant virus, while ART or the immune system clears infected cells. Depending on the precise timing of these treatments, successful approach can be obtained. In this work, we propose two main strategies. The first strategy focuses on simultaneous ART and Shock, where ART and LRAs are applied together. The second strategy use opposite Phase ART and Shock. Here LRAs are given during ART interruption.

In the simultaneous ART and shock approach case, LRAs reactivate latent cells while ART prevents new infections. This strategy reduces viral rebound and keeps viremia low. However, it may limit immune clearance since ART suppresses viral antigen presentation. ART might also weaken LRAs and this can reduce their effectiveness.

The opposite phase ART and shock approach allows viral replication by pausing ART during LRA treatment. This enhances antigen presentation and this potentially improves immune clearance. However, this might cause uncontrolled viral replication, immune escape, and reseeding of latent reservoirs. Its success also depends on how fast reactivated cells can be cleared before they can establish new reservoirs.

In this case the latent cell dynamics is governed by

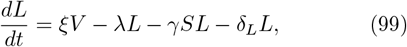

where *ξV*, *λL* and *γSL* denote the formation of new latent cells from infected cells, the reactivation events and the activation due to the latency reversal agent, respectively. *δ*_*L*_*L* represents the natural decay of latent cells.

The infected cell population is given as

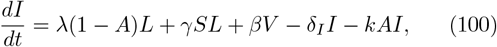

where *λ*(1 − *A*)*L* + *γSL* represents the contribution of reactivated latent cells. *βV* denotes the infection of new CD4+ T cells by free virus while *δ*_*I*_ *I* captures the clearance of infected cells through ART reinitiation. *kAI* accounts for antiviral-induced killing of infected cells. The viral load is governed by:

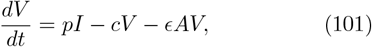

where *pI, cV* and *ϵAV* denote viral production from infected cells, natural viral clearance, and ART-mediated suppression of viral replication, respectively. To incorporate drug washout effects, we introduce the following equations for ART and LRA decay as

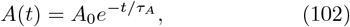

with washout time *τ*_*A*_, and

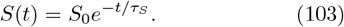

These equation describe the exponential decay of ART and LRA over time after their administration is stopped. Here *τ*_*S*_ represents the washout time of the LRA.

In order to study latent cells, viral load, and infected cells depend on different model parameters, we conducted numerical simulations. Figure 12 depicts that the time evolution of HIV dynamics under ART and LRA treatment. The figure shows that ART (*A* = 0.8) suppresses viral replication and lowers infected cell counts while LRA (*S* = 0.2) activates latent cells by temporarily increasing viral load. As a result, the latent cell population declines steadily due to ART and LRA effects. On the other hand, in Fig, 13 we examine how LRA levels (*S*)influence viral load (*V*). Higher *S* enhances latent cell activation by initially raising *V*. However, excessive activation risks viral rebound before ART can fully suppress replication. Overall, increasing *S* helps clear infected cells and this leads to a decline in *V*. The effect of ART levels (*A*) on viral load is examined in Fig. 14. The figure exhibits that strong ART reduces *V* and this lowers the risk of rebound and reservoir reseeding. However, insufficient ART leads to viral persistence, emphasizing the need for sustained treatment. In Figure 14, we plot LRA levels (*S*) as a function of time for different latent cells (*L*) reservoir. The figure shows that when *S* increases, more latent cells activate and deplete. Excessive activation may trigger viral replication which indicates the need to have critical balance between ART and LRA in optimizing the shock-and-kill strategy.

**FIG. 12:**
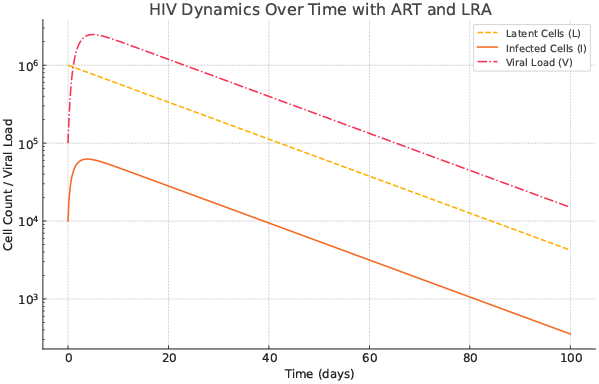
Time evolution of HIV dynamics during ART and LRA treatment. The plot shows the dynamics of latent cells (*L*), infected cells (*I*), and viral load (*V*) as a function of time. The figure shows that ART (*A* = 0.8) suppresses viral replication and infected cells, while LRA (*S* = 0.2) induces reactivation of latent cells. The viral load (*V*) exhibits an initial decline due to ART but shows transient increases due to LRA-induced reactivation. We fix the following parameters as *V*_0_ = 10^5^, *L*_0_ = 10^6^, *I*_0_ = 10^4^, *λ* = 0.05, *γ* = 0.02, *δ*_*L*_ = 0.001, *δ*_*I*_ = 0.5, *β* = 0.001, *p* = 50, *c* = 1, *ϵ* = 0.3, *k* = 0.3.

**FIG. 13:**
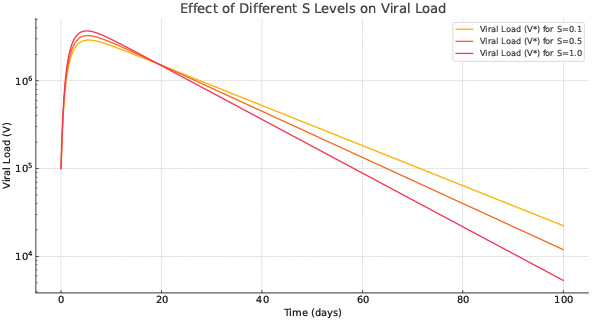
Effect of different LRA levels (*S*) as a function of time and viral load (*V*). An increase in *S* leads to more latent cell activation and this leads to transient increase in viral load. If *S* is too high, the viral load rebounds significantly before ART can suppress it. We fix the parameters as *V*_0_ = 10^5^, *L*_0_ = 10^6^, *I*_0_ = 10^4^, *λ* = 0.05, *γ* = 0.02, *δ*_*L*_ = 0.001, *δ*_*I*_ = 0.5, *β* = 0.001, *p* = 50, *c* = 1, *ϵ* = 0.3, *k* = 0.3.

**FIG. 14:**
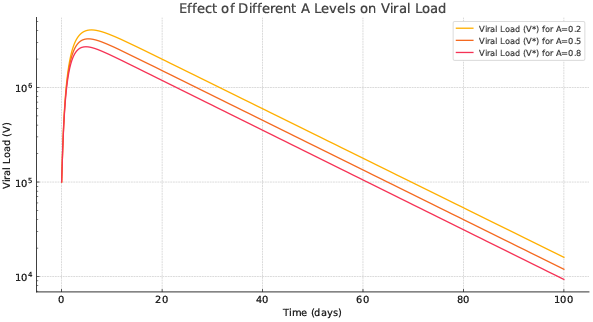
Effect of different ART levels (*A*) on viral load (*V*) over time. the figure depicts that higher ART suppresses viral load more effectively by reducing the risk of viral rebound. If ART is too low, viral load remains elevated, increasing the risk of reservoir reseeding. In the figure, we fix *V*_0_ = 10^5^, *L*_0_ = 10^6^, *I*_0_ = 10^4^, *λ* = 0.05, *γ* = 0.02, *δ*_*L*_ = 0.001, *δ*_*I*_ = 0.5, *β* = 0.001, *p* = 50, *c* = 1, *ϵ* = 0.3, *k* = 0.3.

**FIG. 15:**
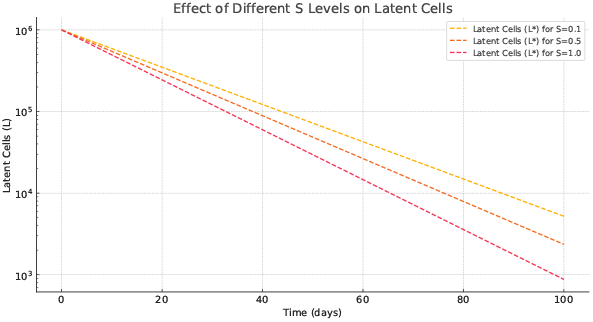
LRA levels (*S*) as a function of time for different latent cells (*L*) reservoir. When *S* increases, it induce more latent cell activation and this leads to a reduction in the latent reservoir. If *S* is too high, a significant number of latent cells transition to active infected cells and this increases viral replication. We plot the figure by fixing *V*_0_ = 10^5^, *L*_0_ = 10^6^, *I*_0_ = 10^4^, *λ* = 0.05, *γ* = 0.02, *δ*_*L*_ = 0.001, *δ*_*I*_ = 0.5, *β* = 0.001, *p* = 50, *c* = 1, *ϵ* = 0.3, *k* = 0.3.

### A. Equilibrium Solutions and Their Implications

To analyze the long-term behavior of the system, we compute the equilibrium solutions (*L, I*, *V* ^∗^) by setting the time derivatives to zero.

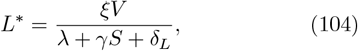

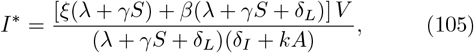

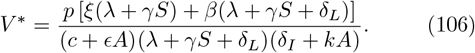

In the limit as *A* → 1, the equilibrium solutions simplify to:

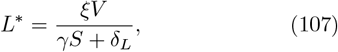

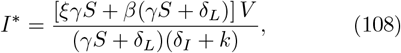

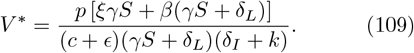

when *A* → 1, ART fully suppresses latent cell activation. This also decreases *L* as well as *I* and *V* because of strong antiviral effects. In the limit *S* → 0, we get

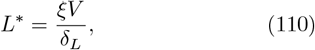

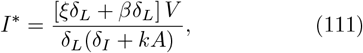

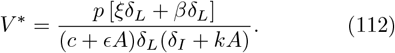

When *S* → 0, latency reversal ceases, leading to a stable but persistent latent reservoir (*L*^∗^ remains higher), while infected cells and viral load decrease as reactivation stops, limiting further infection but preventing complete eradication.

The dependence of latent cells (*L*) as a function of ART level (*A*) and LRA level (*S*) is shown in Fig. 16. As shown in the figure, when *S* increases, it promotes latent cell activation and this decreases *L*. The figure also shows that *A* has minimal direct impact. This indicates that the shock effect from LRA significantly decreases the latent reservoir. Even though, ART does not directly influence latency but it plays a role in limiting new infections. Figure 17 depicts the plot of infected cells (*I*) as a function of ART level (*A*) and LRA level (*S*). When *S* increases, latent cell activation steps up and this increases *I*. However, increasing in *A* suppresses *I* (through antiviral effects). This demonstrates the opposing effects of LRAs and ART. The LRAs induce reactivation and

**FIG. 16:**
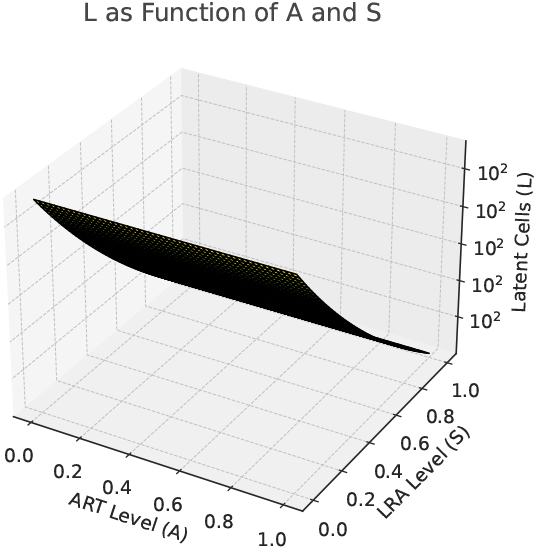
Plot of latent cells (*L*) as a function of ART level (*A*) and LRA level (*S*). The figure depicts that *S* promotes latent cell activation and as a result *L* decreases. As shown in the figure, *A* has minimal direct impact. In the figure we fix *V*_0_ = 10^5^, *L*_0_ = 10^6^, *I*_0_ = 10^4^, *λ* = 0.05, *γ* = 0.02, *δ*_*L*_ = 0.001, *δ*_*I*_ = 0.5, *β* = 0.001, *p* = 50, *c* = 1, *ϵ* = 0.3, *k* = 0.3.

**FIG. 17:**
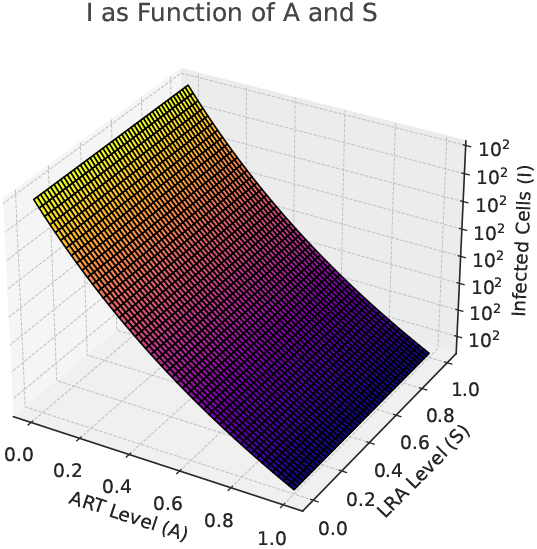
Plot of infected cells (*I*) versus ART level (*A*) and LRA level (*S*). When *S* increases latent cell activation, *I* increases. Because of antiviral effects, increasing in *A* suppresses *I*. The figure is plotted by fixing *V*_0_ = 10^5^, *L*_0_ = 10^6^, *I*_0_ = 10^4^, *λ* = 0.05, *γ* = 0.02, *δ*_*L*_ = 0.001, *δ*_*I*_ = 0.5, *β* = 0.001, *p* = 50, *c* = 1, *ϵ* = 0.3, *k* = 0.3.

ART effectively eliminates newly infected cells. The balance between these two factors is crucial in minimizing viral persistence. In Figure 18, we explore the dependence of (*V*) on ART level (*A*) and LRA level (*S*). Once again as *A* steps up, it effectively suppresses *V*. Higher *S* initially increases viral replication due to latent cell activation.

**FIG. 18:**
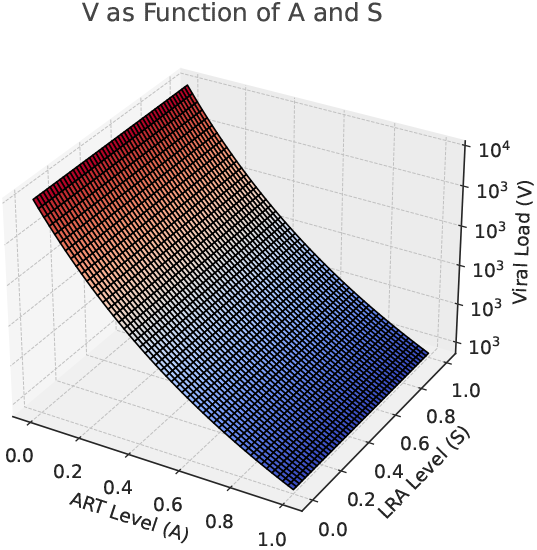
The dependence of (*V*) on ART level (*A*) and LRA level (*S*) is shown in this figure. As shown in the figure *A* effectively suppresses *V*. Due to latent cell activation, higher *S* initially increases viral replication. The parameters are fixed as *V*_0_ = 10^5^, *L*_0_ = 10^6^, *I*_0_ = 10^4^, *λ* = 0.05, *γ* = 0.02, *δ*_*L*_ = 0.001, *δ*_*I*_ = 0.5, *β* = 0.001, *p* = 50, *c* = 1, *ϵ* = 0.3, *k* = 0.3.

This result shows that an important balance is need for an effective treatment. One can note that LRAs activate the virus in the shock phase, but the virus can rebound without enough ART. To prevent this, ART and LRAs must be carefully combined to block viral escape while effectively clearing latent infections. In other words, all of these results stress for the need for a well-planned shock-and-kill strategy since LRAs help reduce the hidden virus by reactivating infected cells and ART is essential to stop the virus from spreading. Finding the right balance between these treatments is key to reducing the latent reservoir and achieving long-term HIV control.

## VII. SUMMARY AND CONCLUSION

In this study, by developing a mathematical framework, we study the stochastic nature of HIV latency reversal and viral reactivation. By modeling reactivation as a Poisson-driven stochastic process, we explore the effect of time-dependent immune fluctuations, ART pharmacokinetics, and random activation events. Our results indicate that the time for viral rebound timing strongly depends on ART decay rates. Slower drug clearance prolongs latency while rapid clearance accelerates reactivation. Furthermore, incorporating Gamma-distributed waiting times revealed that early post-ART activation events are less frequent, while later ones become increasingly probable.

We systematically examined four distinct activation rate scenarios: constant, sinusoidal, randomly fluctuating, and exponentially decreasing rates. The results indicate that immune-driven oscillations lead to clustered reactivation events during peak immune activity, whereas suppression phases prolong latency. Strong stochastic fluctuations significantly increase the risk of viral rebound, while slow immune-driven variations reduce reactivation likelihood. Moreover, our result suggests that early ART suppression limits initial viral expansion. Once drug levels decline, viral replication accelerates exponentially.

The results obtained in this work have direct implications for treatment strategies such as the *Shock-and-Kill* approach. We show that the effectiveness of this strategy improves when the *shock* phase (induction of viral activation by LRAs) is properly synchronized with immune stimulation. In the *kill* phase, proper timing enhances viral clearance which may decrease the risk of reestablishing reservoirs. Our study also depicts that ART decay rates also play a crucial role. As exhibited in this work, slower decay provides a longer window for immunemediated clearance. On the other hand, rapid clearance increases the risk of uncontrolled viral expansion.

In conclusion, our results stress the need for right timing in HIV eradication strategies. Aligning latency-reversing interventions with periods of heightened immune activity could not only enhance treatment efficacy but also improve post-treatment viral control. One should note that also addressing the stochastic nature of reactivation remains a key challenge since this requires a combination of mathematical modeling, immunological insights, and precise medicine approaches to optimize long-term remission strategies.

## Acknowledgment

I would like to thank Mulu Zebene for the constant encouragement.

## Data Availability Statement

This manuscript has no associated data or the data will not be deposited. [Authors’ comment: Since we presented an analytical work, we did not collect any data from simulations or experimental observations.]

## Notes

### Competing Interest Statement

The authors have declared no competing interest.

